# Implications of *de novo* mutations in guiding drug discovery: A study of four neuropsychiatric disorders

**DOI:** 10.1101/173641

**Authors:** Hon-Cheong So, Yui-Hang Wong

## Abstract

Recent studies have suggested an important role of *de novo* mutations (DNMs) in neuropsychiatric disorders. As DNMs are not subject to elimination due to evolutionary pressure, they are likely to have greater disruptions on biological functions. While a number of sequencing studies have been performed on neuropsychiatric disorders, the implications of DNMs for drug discovery remain to be explored.

In this study, we employed a gene-set analysis approach to address this issue. Four neuropsychiatric disorders were studied, including schizophrenia (SCZ), autistic spectrum disorders (ASD), intellectual disability (ID) and epilepsy. We first identified gene-sets associated with different drugs, and analyzed whether the gene-set pertaining to *each* drug overlaps with DNMs more than expected by chance. We also assessed which medication classes are enriched among the prioritized drugs. We discovered that neuropsychiatric drug classes were indeed significantly enriched for DNMs of all four disorders; in particular, antipsychotics and antiepileptics were the *most* strongly enriched drug classes for SCZ and epilepsy respectively. Interestingly, we revealed enrichment of several unexpected drug classes, such as lipid-lowering agents for SCZ and anti-neoplastic agents. By inspecting individual hits, we also uncovered other interesting drug candidates or mechanisms (*e.g.* histone deacetylase inhibition and retinoid signaling) that might warrant further investigations. Taken together, this study provided evidence for the usefulness of DNMs in guiding drug discovery or repositioning.

## Introduction

The past decade has witnessed rapid development in sequencing technologies. Whole-exome and whole-genome sequencing enables the discovery of many *de novo* mutations (DNM) (mutations present in the offspring but absent in either parent) in Mendelian as well as complex diseases. Recent studies have suggested an important role of DNM in neuropsychiatric disorders, such as schizophrenia (SCZ), autistic spectrum disorders (ASD), intellectual disability (ID) and epilepsy^1-3^. De novo mutations are rare and unlike inherited variants, they are not subject to elimination due to evolutionary pressure. They are therefore likely to be have larger effect sizes on disease risk and more significant disruptions on biological functions^3^. While a relatively large number of sequencing studies have been performed on neuropsychiatric disorders, their implications for the development for new therapies are rarely explored. Ideally, for affected individuals harboring DNM that are likely pathogenic, a “precision medicine” approach can be applied, such that the therapy will specifically target the mutations. This approach is however challenging and costly as hundreds of mutations have been identified for each of the abovementioned neuropsychiatric disorders.

In this study, we investigated another approach by considering the collection of DNM instead of focusing on a single mutation. We hypothesized that the *DNM* as a whole will reflect disease pathophysiology, and they might be associated with drugs known to treat or potentially useful for the diseases under study.

We focused on four neuropsychiatric disorders (SCZ, ASD, ID, epilepsy) here. Recent studies have shown genetic overlap between these four disorders^4^ and hence we will study them together. In terms of pharmacological treatment, a number of antipsychotics and antiepileptics have been developed for SCZ and epilepsy respectively. However, as a whole, different psychiatric medications are also commonly prescribed for these disorders, including for ASD and ID^5-8^.

We are interested in this question: are gene-sets associated with neuropsychiatric drug classes over-represented among the DNM? Specifically, we hypothesized that antipsychotic gene-sets might be over-represented among the SCZ *de novo* mutations, and a similar relationship exists for antiepileptics and epilepsy mutations. We also expect enrichment of neuropsychiatric drug classes for ASD and ID due to shared genetic bases^4,9^ and clinical comorbidities^10,11^ with other neuropsychiatric diseases. If our hypothesis is true, the approach also serves as a way for drug discovery or repositioning based on *DNM*: drugs that are significantly over-represented (but not indicated for the disorder) can serve as candidates for repositioning. It is worth mentioning that here we adopted a “multi-target” paradigm for drug discovery. While conventional drug development focus on single drug targets, many diseases involve complex interplay of multiple genes and pathways, and it has been argued that a multi-target approach may reveal drugs with better efficacies^12-14^. Indeed many drugs that are effective and widely used in treating neuropsychiatric disorders are multi-target, such as valproate^15^ and clozapine^16^.

Our analyses can be broadly divided into two steps. Firstly, we identified gene-sets related to a variety of drugs, and analyzed whether the gene-set pertaining to *each* drug overlaps with DNM more than expected by chance. This is very similar to a pathway analysis performed on a set of candidate genes. A gene-set related to a drug can be viewed as a “pathway” in this case. The top-ranked drugs may then serve as repositioning candidates. Secondly, we further analyzed the prioritized drugs, and assessed which drug classes were enriched among the top results. As discussed above, we will test specifically if several neuropsychiatric drug classes are enriched.

## Methods

### *De novo* mutation resources

We made use of two recently developed databases, NPdenovo^4^ and denovo-db^17^, for the current analyses. NPdenovo (www.wzgenomics.cn/NPdenovo/) is a database dedicated to neurodevelopmental disorders, including SCZ, ASD, ID and epileptic encephalopathy (EE). Details of database construction and curation can be found in Li et al.^4^. Briefly, information of 3555 trios for the 4 aforementioned disorders together with unaffected siblings or controls were collected from whole-exome sequencing (WES) or whole-genome sequencing (WGS) studies. In total 17104 DNM were extracted and annotated using various bioinformatics tools. NPdenovo also categorized some mutations with high estimated pathogenicity as “extreme” mutations. Several kinds of likely gene-disrupting (LGD) events such as nonsense, splice-site and frameshift mutations are directly considered to be damaging or extreme. A large proportion of DNMs are however missense mutations, for which pathogenicity are less clear. The NPdenovo database integrates functional predictions of the damaging ability of each variant from 12 computational tools. An aggregate “damaging score” is computed by summing up the number of tools that predict the variant to be deleterious, and any variant with a score >= 8 is considered an “extreme” mutation. Details of this procedure are described in Li et al^4^. We also considered another database denovo-db that is slightly more recent; it is a more comprehensive resource covering all kinds of disorders. The database provides the Combined Annotation Dependent Depletion (CADD) score^18^ as a measure of likely pathogenicity, but did not categorize variant directly into “extreme” or “non-extreme” mutations. Readers are referred to Turner et al.^17^ for details of the denovo-db.

### Gene-set associated with each drug

We made use of the DSigDB database^19^ to extract gene-sets related to each drug. The whole database was downloaded from http://tanlab.ucdenver.edu/DSigDB. A total of 17839 unique compounds were included in this database. The gene-sets were compiled according to multiple sources: (1) bioassay results from PubChem^20^ and ChEMBL^21^; (2) kinase profiling assay from the literature and two kinase databases; (3) drug-induced differentially expressed genes (with >2 fold-change compared to controls), as derived from the Connectivity Map^22^; and (4) manually curated and text mined drug targets from the Therapeutics Targets Database^23^ and the Comparative Toxicogenomics Database^24^.

### Testing for overlap of *de novo* mutations with drug-related genes

We then tested for over-representation of drug-related gene-sets among the DNMs. This is very similar to a pathway analysis performed on candidate genes, but here the “pathway” is a set of genes related to the drug action. We constructed for each drug a 2 x 2 contingency table, summarizing the total number of genes categorized by: (1) whether or not the gene belongs to a (specified type of) DNM; and (2) whether or not the gene is associated with the particular drug.

We tested for independence of the two categories by a one-tailed Fisher’s exact test (we were testing for a *greater*-than-expected overlap, hence one-tailed tests). We hypothesized that some drugs will target genes that overlap with DNM more than expected by chance; these drugs can be prioritized as repositioning candidates. To avoid results driven by too few genes and since we are focusing on a “multi-target” paradigm of drug discovery, we excluded drugs with less than 5 associated genes. A total of 5389 drugs were included in the final analyses.

### Stratification of de novo mutations

As the functional significance and pathogenicity of DNMs may differ a lot, we performed analyses with different subgroups of DNMs. For mutations in NPdenvo, we repeated the over-representation analysis with all non-synonymous and then “extreme” mutations. For denovo-db, we first included mutations which are LGD (including stop-gained, stop-loss, frameshift, splice acceptor or donor mutations) as pathogenic variants. We then stratified missense mutations by their CADD scores. It should be noted that there is no consensus or strong theoretical basis for a particular cutoff of the score. Here we followed the suggestion by the authors (http://cadd.gs.washington.edu/info) and set 15 as a primary cutoff for pathogenic mutations; we also performed a subsidiary analyses with a much more stringent threshold of 30. In addition, for both databases, we stratified the mutations according to whether they are exclusively found in cases or also found in control subjects.

### Testing for drug class enrichment

After we prioritized the drugs from the previous step, we tested for enrichment of individual drug classes. Briefly, we tested whether drugs of a particular class (e.g. anti-epileptics or antipsychotics) would have significantly lower p-values than drugs not belonging to that class. P-values were converted to z-statistics by a probit function [*z* = Φ^−1^ (*p*)] and one-tailed t-tests were used to compare the means of drugs within or outside the specified drug class. (This is analogous to the principle of gene-set analysis described in ref.^25,26^.)

As discussed earlier, we are specifically interested in the enrichment of neuropsychiatric drug classes, as they are most pertinent to the diseases under study. We specifically extracted drugs belonging to (1) antipsychotics; (2) antidepressants and anxiolytics; (3) antiepileptics; and (4) all psychiatric drugs, which include drugs for schizophrenia, bipolar disorder, depression, anxiety and phobic disorders, ASD, attention deficit hyperactivity disorder and dementia (including Alzheimer’s disease). Drug classes were defined based on two sources, including the Anatomical Therapeutic Chemical (ATC) Classification System and the MEDI-HPS (MEDication Indication – High Precision Subset)^27^. ATC is a classification system set by the World Health Organization for therapeutic agents, while MEDI is an ensemble resource of medication indication formed by integrating four commonly used medication resources (RxNorm, MedlinePlus, SIDER2^28^ and Wikipedia). MEDI-HPS is a subset of MEDI which only includes indications found in either RxNorm or at least 2 of the 3 other sources, with an estimated precision of 92% based on clinician review^27^.

As will be discussed in the results section, we observed highly significant enrichment of neuropsychiatric drug classes for NPdenovo “extreme” mutations that are limited to cases. For this group of mutations, we also performed a more comprehensive enrichment test covering *all* level 3 ATC drug classes; the purpose is to examine drug classes (in addition to individual drugs) that may have potential for repositioning. Similar to our previous analyses, in order to avoid significant results driven by few drugs in a category, we included pharmacological classes with at least 5 drugs. A total of 128 pharmacological classes were included for the final analysis.

### Literature search of prioritized drugs

For drugs prioritized by the analysis of NPdenovo extreme mutations (exclusive in cases), we also performed a systemic literature search in PubMed and Google Scholar for possible therapeutic relevance. The search strategy is as follows: (1) For SCZ: Drug_name AND (schizophrenia OR schizophrenic OR psychosis OR psychotic OR antipsychotic); (2) for ASD: Drug_name AND (autism OR autistic OR asperger); (3) for ID: Drug_name AND (“general learning disability” OR “intellectual disability” OR “mental retardation”); (4) for epilepsy: Drug_name AND (epilepsy OR epileptic OR seizure). Clinical studies were cited in preference to pre-clinical studies if available. Systematic review and meta-analysis were cited with higher priorities, generally following the hierarchy of evidence. Annotation was performed for the top 30 drugs for each disorder. A maximum of 3 references are included. The literature search was performed in Jul 2017. Due to the relatively large number of drug-disorder pairs, this is not meant to be a systematic review of current evidence but an approximate guide to the therapeutic relevance of each top-ranked drug, providing a reference for other researchers.

### Multiple testing correction

We employed the false discovery rate (FDR) approach for multiple testing correction, which controls the expected proportion of false discoveries^29^. FDR correction was performed by the Benjamini-Hochberg procedure implemented in the R function p.adjust. The corresponding “adjusted p-values”, or q-values^30^, were reported. Results with q-values less than 0.05 were regarded as significant associations, while results with 0.05 < *q* <=0.1 were considered suggestive associations.

## RESULTS

### Drug class enrichment results from NPdenovo

Table 1 shows the enrichment of neuropsychiatric drug classes for SCZ and ASD from analysis of the NPdenovo database. Antipsychotics were strongly enriched for SCZ DNM (lowest *p* = 4.76E-9 from four analyses of different subtypes of DNMs). The enrichment was highly significant regardless of the subtype of DNM we analyzed. Nevertheless, the enrichment was stronger for extreme DNMs, and also slightly stronger for DNMs found exclusively in cases. Interestingly, there was also enrichment for antiepileptics (lowest *p* = 1.69E-4), and as expected, for all psychiatric drugs (lowest *p* = 1.31E-5). There was a modest association with medications for depression and anxiety defined by MEDI-HPS (lowest *p* = 2.37E-2). For ASD, the strongest enrichment was for antiepileptics (lowest *p* = 3.01E-11), but we also detected an enrichment for MEDI-HPS drugs for depression and anxiety (lowest *p* = 4.17E-4), and for psychiatric medications in general.

**Table 1.** Enrichment of neuropsychiatric drug classes based on *de novo* mutations in NPdenovo database (SCZ and ASD)

|  | SCZ non-syn | SCZ extreme | ASD non-syn | ASD extreme |
| --- | --- | --- | --- | --- |
| *ATC classification* |  |  |  |  |
| Antipsychotics | **6.59E-06** | **1.33E-07** | 9.43E-01 | 8.21E-01 |
| Antidepressants or anxiolytics | 8.37E-01 | 6.16E-01 | 1.76E-01 | 8.15E-01 |
| All ATC psychiatric drugs | 3.38E-01 | **2.58E-02** | **2.46E-02** | *3.33E-02* |
| Antiepileptics | **1.38E-03** | **6.50E-04** | **3.50E-08** | **1.93E-08** |
| *MEDI-HPS* |  |  |  |  |
| For Schizophrenia and bipolar | **6.17E-05** | **9.60E-06** | 4.50E-01 | 2.48E-01 |
| For MDD or anxiety disorders | 4.10E-01 | 7.01E-02 | **9.64E-04** | **1.39E-02** |
| All psychiatric drugs | **9.50E-03** | **4.74E-04** | **8.59E-03** | **1.65E-02** |
| Antiepileptics | **2.42E-02** | **4.84E-03** | **1.23E-09** | **4.33E-11** |
| *Limited to mutations exclusively found in cases* |  |  |  |  |
| *ATC classification* |  |  |  |  |
| Antipsychotics | **2.68E-06** | **4.76E-09** | 8.86E-01 | 8.77E-01 |
| Antidepressants or anxiolytics | 6.24E-01 | 3.37E-01 | 1.00E-01 | 7.12E-01 |
| All ATC psychiatric drugs | 1.10E-01 | **1.10E-03** | **1.30E-02** | **3.39E-02** |
| Antiepileptics | **1.70E-04** | **1.69E-04** | **1.68E-06** | **1.99E-08** |
| *MEDI-HPS* |  |  |  |  |
| For Schizophrenia and bipolar | **1.53E-05** | **2.93E-07** | 3.37E-01 | 2.27E-01 |
| For MDD or anxiety disorders | 1.76E-01 | **2.37E-02** | **4.17E-04** | **7.36E-03** |
| All psychiatric drugs | **8.23E-04** | **1.31E-05** | **5.81E-03** | **1.11E-02** |
| Antiepileptics | **7.87E-03** | **3.52E-03** | **4.10E-08** | **3.01E-11** |

SCZ, schizophrenia; ASD, autistic spectrum disorder.
Non-syn, all non-synonymous DNMs; extreme, extreme mutations as defined in the NPdenovo database.
Results with q-values $\leq 0.05$ are in bold, and those with $0.05 < q \leq 0.1$ are in italics.

Table 2 shows the enrichment of neuropsychiatric drug classes for ID and epilepsy from NPdenovo. For ID, a strong enrichment was observed for antiepileptics (lowest p = 2.67E-8). There was also evidence of enrichment for antipsychotics and antidepressants/anxiolytics, no matter the drug classes were defined by ATC or MEDI-HPS. Finally, for epilepsy, the most significant enrichment was for antiepileptics (lowest p = 8.69E-12), but there was also evidence that drugs for depression and anxiety were enriched (lowest p = 5.20E-3). Note that all the above-mentioned associations were significant at an FDR threshold of 0.05.

**Table 2.**
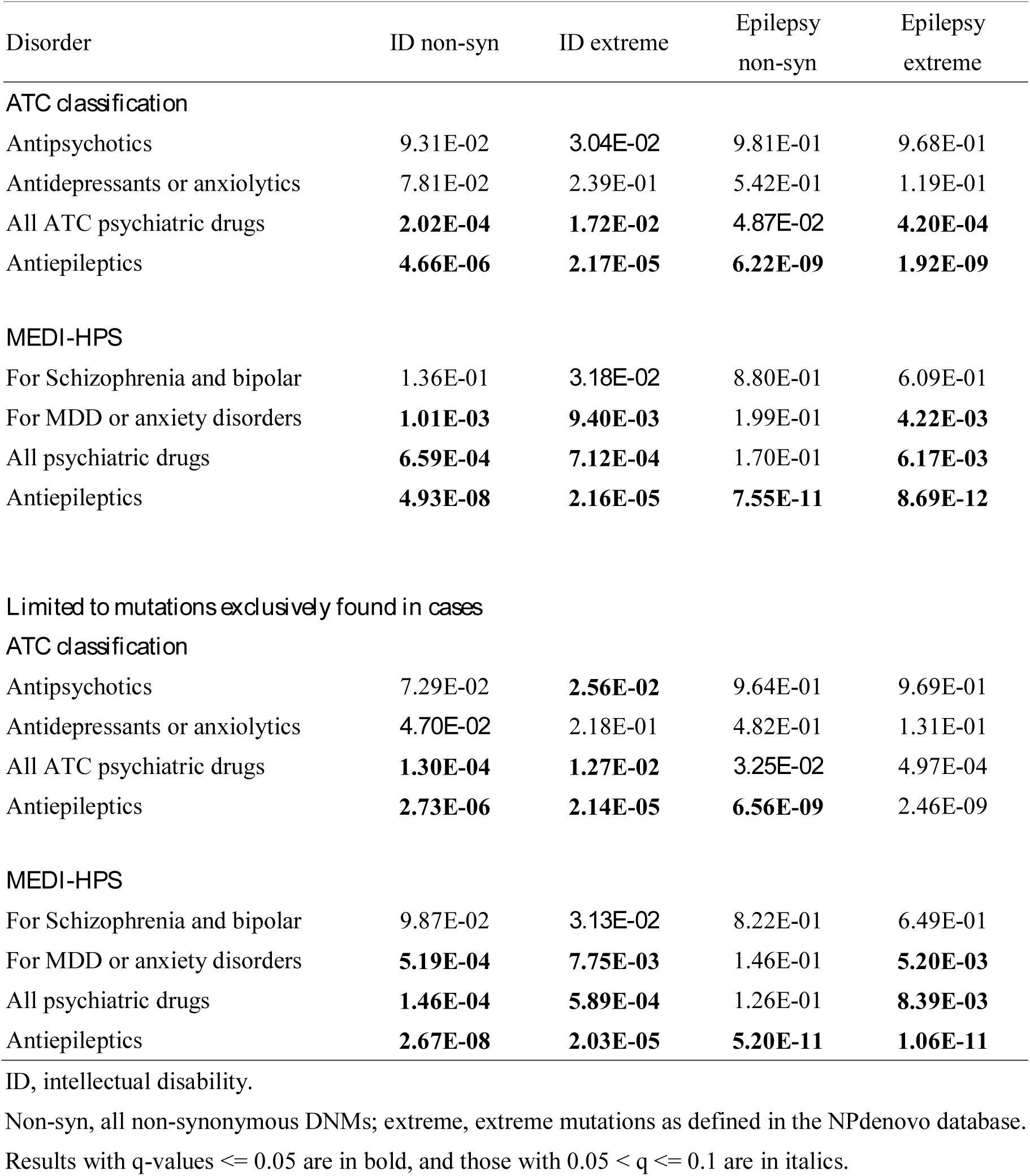
Enrichment of neuropsychiatric drug classes based on *de novo* mutations in NPdenovo database (ID and epilepsy)

For all four neuropsychiatric disorders under study, there was significant enrichment for the set of combined psychiatric medications. For SCZ and epilepsy, the two disorders with well-established treatments, the strength of enrichment (*e.g.* for antipsychotics and antiepileptics) tended to be stronger when we focused on “extreme” mutations and mutations exclusively detected in cases. For ID and ASD, the difference was not obvious. One possible explanation is that ID and ASD have no established drug treatments; the therapeutic relevance of psychiatric medications may depend on the specific (comorbid) clinical symptoms of patients, instead of the severity of the mutations causing the disorder. For example, a patient with ID may have a DNM in a gene that also predisposes to depressive symptoms; antidepressants may be relevant to this patient, even if the DNM is not highly pathogenic for ID. On the other hand, more deleterious or “extreme” mutations for SCZ or epilepsy may indicate a more important role in disease pathogenesis, and hence higher potential as therapeutic targets.

### Drug class enrichment results from denovo-db

The enrichment test results from denovo-db were broadly similar to those obtained from analysis of NPdenovo. Results of analysis were shown in Table 3-4. Again we observed strong enrichment for antipsychotics in SCZ and antiepileptics in epilepsy. We also observed enrichment for medications against depression and anxiety (from MEDI-HPS) across all four disorders. Antiepileptics and the combined drug-set of psychiatric medications were also enriched across all four disorders. The enrichment appeared to be roughly similar for mutations limited to cases or not, although there was sometimes a slightly stronger enrichment for mutations exclusive in cases. The significance of enrichment was also not much different for the two CADD cutoffs, except for SCZ. Interestingly, in SCZ, enrichment for neuropsychiatric drug classes was no longer significant when we increased the CADD score threshold to 30, except for antiepileptics. One explanation is that there are fewer variants passing a threshold score of 30, which impairs the power of detecting associations; in addition, setting a very stringent CADD threshold at 30 may have stratified the SCZ patients into a distinct subgroup that is different from the more typical patients. It is worth noting that not all SCZ patients have DNMs and common genetic variants played a significant role in disease pathogenesis^31^.

**Table 3.** Enrichment of neuropsychiatric drug classes based on *de novo* mutations listed in denovo-db (SCZ and ASD)

| Disorder | SCZ_cadd15 | SCZ_cadd30 | ASD_cadd15 | ASD_cadd30 |
| --- | --- | --- | --- | --- |
| *ATC classification* |  |  |  |  |
| Antipsychotics | **1.11E-07** | 1.38E-01 | 9.93E-01 | 8.49E-01 |
| Antidepressants or anxiolytics | 1.02E-01 | 9.73E-01 | 7.39E-01 | 8.05E-01 |
| All ATC psychiatric drugs | **4.59E-03** | 7.80E-01 | 2.09E-01 | 5.19E-01 |
| Antiepileptics | **4.65E-04** | **8.89E-04** | **2.65E-09** | **2.87E-10** |
| *MEDI-HPS* |  |  |  |  |
| For Schizophrenia and bipolar | **7.42E-08** | 9.23E-02 | 8.88E-01 | 6.58E-01 |
| For MDD or anxiety disorders | **1.99E-03** | 1.36E-01 | 8.91E-02 | **1.19E-02** |
| All psychiatric drugs | **5.10E-07** | 1.01E-01 | 1.85E-01 | 2.36E-01 |
| Antiepileptics | **3.44E-03** | **1.86E-03** | **1.71E-07** | **2.55E-12** |
| *Limited to mutations exclusively found in cases* |  |  |  |  |
| *ATC classification* |  |  |  |  |
| Antipsychotics | **1.28E-07** | 1.42E-01 | 9.92E-01 | 8.94E-01 |
| Antidepressants or anxiolytics | 5.62E-02 | 9.72E-01 | 7.47E-01 | 5.31E-01 |
| All ATC psychiatric drugs | **1.10E-03** | 7.88E-01 | 3.08E-01 | 3.49E-01 |
| Antiepileptics | **1.55E-04** | **8.79E-04** | **2.02E-09** | **5.03E-11** |
| *MEDI-HPS* |  |  |  |  |
| For Schizophrenia and bipolar | **2.21E-08** | 9.02E-02 | 8.91E-01 | 5.82E-01 |
| For MDD or anxiety disorders | **6.34E-04** | 1.34E-01 | 8.51E-02 | **1.18E-03** |
| All psychiatric drugs | **4.79E-08** | 9.82E-02 | 1.76E-01 | 1.27E-01 |
| Antiepileptics | **1.17E-03** | **1.84E-03** | **1.86E-07** | **1.76E-13** |

SCZ, schizophrenia; ASD, autistic spectrum disorder.
Cadd15, DNMs with CADD score > 15; Cadd30, DNMs with CADD score > 30. Note that we also included all likely gene disrupting (LGD) mutations, including stop-gained, stop-loss, frameshift, splice acceptor or donor mutations.
Results with q-values <= 0.05 are in bold, and those with 0.05 < q <= 0.1 are in italics.

**Table 4.** Enrichment of neuropsychiatric drug classes based on *de novo* mutations listed in denovo-db (ID and epilepsy)

|  | ID_cadd15 | ID_cadd30 | Epilepsy_cadd15 | Epilepsy_cadd30 |
| --- | --- | --- | --- | --- |
| *ATC classification* |  |  |  |  |
| Antipsychotics | 6.52E-01 | 2.22E-01 | 9.44E-01 | 7.89E-01 |
| Antidepressants or anxiolytics | 1.51E-01 | 7.33E-01 | 5.62E-01 | 2.58E-01 |
| All ATC psychiatric drugs | **2.46E-02** | *5.23E-02* | **6.91E-03** | **5.77E-03** |
| Antiepileptics | **9.25E-09** | **1.05E-07** | **2.34E-10** | **6.11E-10** |
| *MEDI-HPS* |  |  |  |  |
| For Schizophrenia and bipolar | 6.53E-01 | 1.21E-01 | 8.71E-01 | 5.55E-01 |
| For MDD or anxiety disorders | **7.44E-04** | **1.18E-02** | **1.33E-02** | **3.36E-03** |
| All psychiatric drugs | **2.14E-02** | **4.88E-03** | 6.31E-02 | **3.65E-03** |
| Antiepileptics | **9.97E-11** | **2.53E-08** | **1.14E-13** | **2.79E-11** |
| *Limited to mutations exclusively found in cases* |  |  |  |  |
| *ATC classification* |  |  |  |  |
| Antipsychotics | 6.50E-01 | 2.35E-01 | 9.41E-01 | 8.79E-01 |
| Antidepressants or anxiolytics | 2.33E-01 | 7.37E-01 | 5.48E-01 | 2.89E-01 |
| All ATC psychiatric drugs | *3.24E-02* | 6.53E-02 | **6.25E-03** | **2.41E-02** |
| Antiepileptics | **7.97E-09** | **9.70E-08** | **2.50E-10** | **6.27E-10** |
| *MEDI-HPS* |  |  |  |  |
| For Schizophrenia and bipolar | 7.32E-01 | 1.33E-01 | 8.54E-01 | 7.35E-01 |
| For MDD or anxiety disorders | **1.51E-03** | **1.12E-02** | **1.26E-02** | **3.41E-03** |
| All psychiatric drugs | *3.07E-02* | **6.40E-03** | *5.75E-02* | **1.71E-02** |
| Antiepileptics | **7.52E-11** | **2.34E-08** | **1.36E-13** | **2.42E-11** |

ID, intellectual disability.
Cadd15, DNMs with CADD score > 15; Cadd30, DNMs with CADD score > 30. Note that we also included all likely gene disrupting (LGD) mutations, including stop-gained, stop-loss, frameshift, splice acceptor or donor mutations.
Results with q-values <= 0.05 are in bold, and those with 0.05 < q <= 0.1 are in italics.

### Enrichment for *all* ATC drug classes for extreme mutations limited to cases

As discussed earlier, we also performed a more comprehensive enrichment test covering all ATC (level 3) drug classes for NPdenovo extreme mutations limited to cases. The results are shown in Table 5. For schizophrenia, the most strongly enriched drug class is antipsychotics, while antiepileptics ranked fifth. Interestingly, lipid modifying agents ranked second with a slightly higher p-value than antipsychotics (*p* = 5.194E-9). Antimicrobial agents including anti-fungal drugs were also ranked high on the list. For ASD, the strongest enrichment was observed for antiepileptics; other groups of psychiatric medications, such as hypnotics/sedatives, anti-dementia drugs and anxiolytics were also ranked among the top with *q*-value < 0.05. Other top-ranked drug classes included anti-neoplastic drugs and antimetabolites, as well as calcium channel blockers. For ID, anti-neoplastic agents and antiepileptics were listed among the top results. As for epilepsy, agents known to treat seizures or raise seizure thresholds such as antiepileptics, hypnotics/sedatives and anxiolytics were all highly enriched.

**Table 5.**
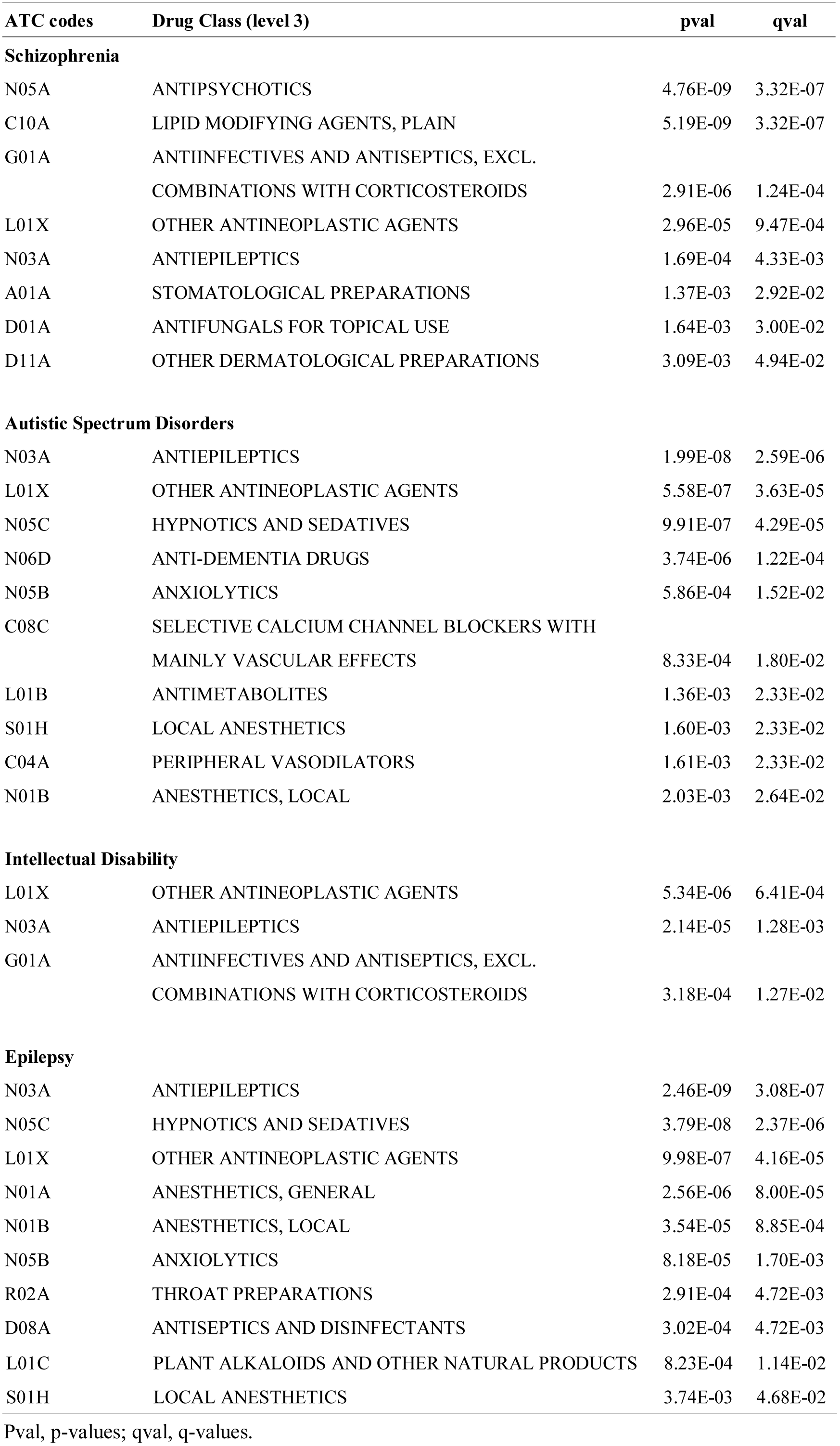
Enrichment of all ATC (level 3) drug classes (only drug classes with q-value < 0.05 are shown)

| ATC codes | Drug Class (level 3) | pval | qval |
| --- | --- | --- | --- |
| **Schizophrenia** |  |  |  |
| N05A | ANTIPSYCHOTICS | 4.76E-09 | 3.32E-07 |
| C10A | LIPID MODIFYING AGENTS, PLAIN | 5.19E-09 | 3.32E-07 |
| G01A | ANTIINFECTIVES AND ANTISEPTICS, EXCL.<br>COMBINATIONS WITH CORTICOSTEROIDS | 2.91E-06 | 1.24E-04 |
| L01X | OTHER ANTINEOPLASTIC AGENTS | 2.96E-05 | 9.47E-04 |
| N03A | ANTIEPILEPTICS | 1.69E-04 | 4.33E-03 |
| A01A | STOMATOLOGICAL PREPARATIONS | 1.37E-03 | 2.92E-02 |
| D01A | ANTIFUNGALS FOR TOPICAL USE | 1.64E-03 | 3.00E-02 |
| D11A | OTHER DERMATOLOGICAL PREPARATIONS | 3.09E-03 | 4.94E-02 |
| **Autistic Spectrum Disorders** |  |  |  |
| N03A | ANTIEPILEPTICS | 1.99E-08 | 2.59E-06 |
| L01X | OTHER ANTINEOPLASTIC AGENTS | 5.58E-07 | 3.63E-05 |
| N05C | HYPNOTICS AND SEDATIVES | 9.91E-07 | 4.29E-05 |
| N06D | ANTI-DEMENTIA DRUGS | 3.74E-06 | 1.22E-04 |
| N05B | ANXIOLYTICS | 5.86E-04 | 1.52E-02 |
| C08C | SELECTIVE CALCIUM CHANNEL BLOCKERS WITH<br>MAINLY VASCULAR EFFECTS | 8.33E-04 | 1.80E-02 |
| L01B | ANTIMETABOLITES | 1.36E-03 | 2.33E-02 |
| S01H | LOCAL ANESTHETICS | 1.60E-03 | 2.33E-02 |
| C04A | PERIPHERAL VASODILATORS | 1.61E-03 | 2.33E-02 |
| N01B | ANESTHETICS, LOCAL | 2.03E-03 | 2.64E-02 |
| **Intellectual Disability** |  |  |  |
| L01X | OTHER ANTINEOPLASTIC AGENTS | 5.34E-06 | 6.41E-04 |
| N03A | ANTIEPILEPTICS | 2.14E-05 | 1.28E-03 |
| G01A | ANTIINFECTIVES AND ANTISEPTICS, EXCL.<br>COMBINATIONS WITH CORTICOSTEROIDS | 3.18E-04 | 1.27E-02 |
| **Epilepsy** |  |  |  |
| N03A | ANTIEPILEPTICS | 2.46E-09 | 3.08E-07 |
| N05C | HYPNOTICS AND SEDATIVES | 3.79E-08 | 2.37E-06 |
| L01X | OTHER ANTINEOPLASTIC AGENTS | 9.98E-07 | 4.16E-05 |
| N01A | ANESTHETICS, GENERAL | 2.56E-06 | 8.00E-05 |
| N01B | ANESTHETICS, LOCAL | 3.54E-05 | 8.85E-04 |
| N05B | ANXIOLYTICS | 8.18E-05 | 1.70E-03 |
| R02A | THROAT PREPARATIONS | 2.91E-04 | 4.72E-03 |
| D08A | ANTISEPTICS AND DISINFECTANTS | 3.02E-04 | 4.72E-03 |
| L01C | PLANT ALKALOIDS AND OTHER NATURAL PRODUCTS | 8.23E-04 | 1.14E-02 |
| S01H | LOCAL ANESTHETICS | 3.74E-03 | 4.68E-02 |

Pval, p-values; qval, q-values.

### Literature search results of the top prioritized drugs

The full results of the literature search are listed in supplementary Table 1-4. Some of the drugs with potential therapeutic relevance are highlighted in Table 6.

**Table 6.**
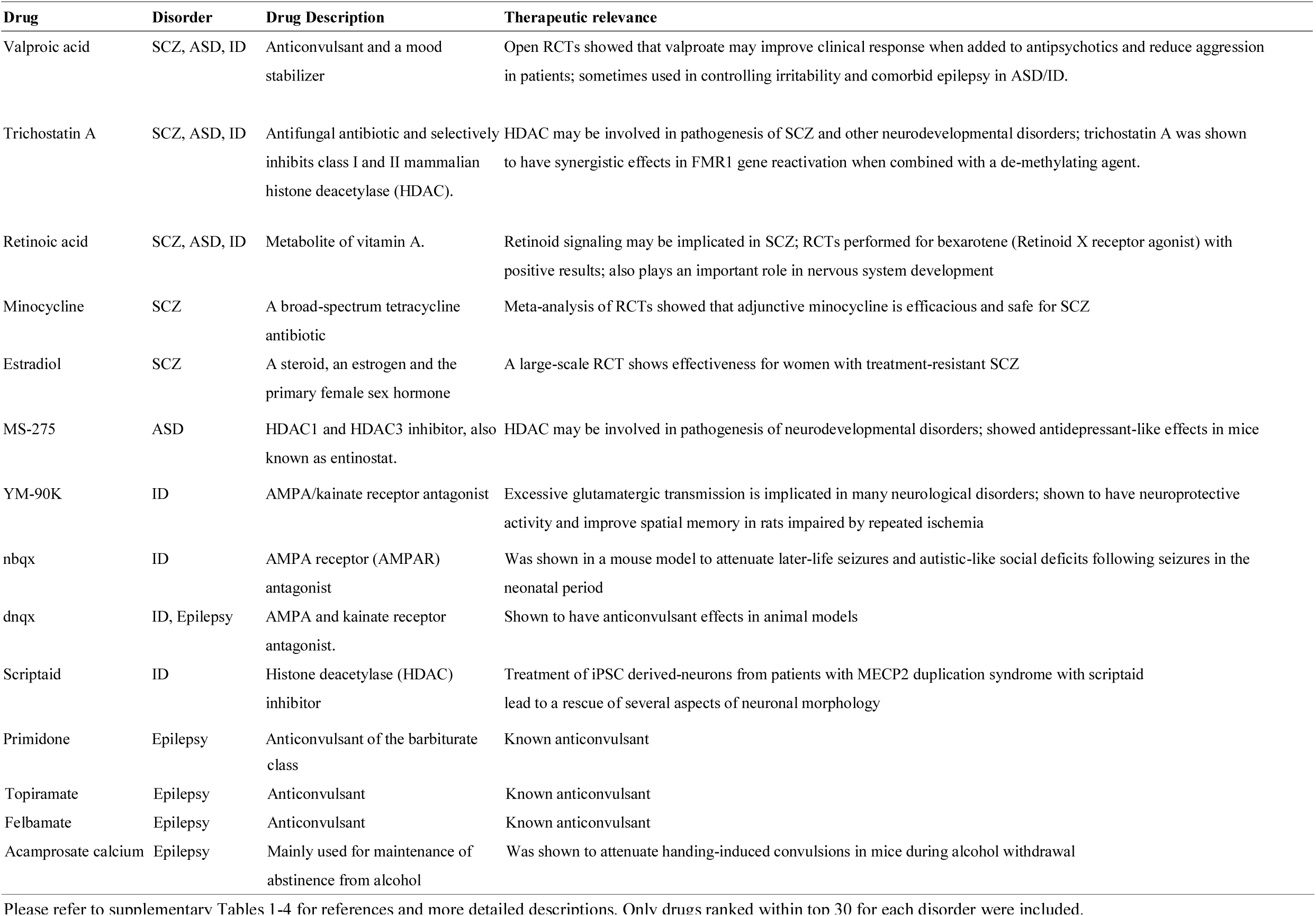
Selected drugs enriched for individual disorders (ranked within top 30) and therapeutic relevance

#### Top-ranked drugs across disorders

A few drugs are listed across disorders and are discussed here. Histone deacetylases inhibitors (HDAC) inhibitors such as trichostatin A, MS-275 and scriptaid were listed among the top for SCZ, ASD and ID. In fact the commonly used mood stabilizer valproate is also an HDAC inhibitor. HDAC is believed to play an important role in many complex diseases, including neuropsychiatric disorders^32,33^. HDAC removes acetyl groups from protein substrates on histones, and may lead to transcriptional silencing by allowing for chromatin compaction; HDAC inhibitors in turn block the deacetylation process^32^. HDAC inhibitors have been suggested as new therapies for neuropsychiatric disorders^32-35^. For instance, HDAC inhibitor has been suggested to as a new therapy against positive, negative and cognitive symptoms in schizophrenia^36^, mainly based on pre-clinical evidence. There is also evidence for HDAC inhibitors in the treatment of neurodevelopmental disorders such as ID. Fragile X syndrome (FXS) is the most common inherited cause of ID and the most common monogenetic cause of ASD^37^. Hypermethylation and increased histone deacetylation of the *FMR* gene has been shown to contribute to the disease^38,39^. Treatment of lymphoblastoid cells derived from FXS patients with 5-aza-2-deoxycytidine (a de-methylating agent) together with HDAC inhibitors was reported to have synergistic effects in reactivating the *FMR1* gene^39^. HDAC inhibitors were also implicated in the treatment of another syndromal cause of ID, Rubinstein-Taybi syndrome^40^. Decreased histone acetylation appears to be a common mechanism underlying many neurodevelopmental or other brain disorders^33^. However, current evidence was largely based on pre-clinical studies and further clinical investigations including randomized controlled trials (RCTs) are necessary to validate these findings.

Another drug, retinoic acid, was also listed among the top across SCZ, ASD and ID. Retinoic acid signaling plays an important part in the development of the central nervous system, for example in neural differentiation, axon outgrowth and neural patterning^41,42^. For schizophrenia, the retinoid signaling pathway was proposed as a novel therapeutic target^43^, and a retinoid X receptor agonist (bexarotene) has been tested in two clinical trials^44,45^ with some evidence of benefit in SCZ as an add-on agent. Retinoic acid was proposed as a potential therapy for ASD as well, based on its action of inducing CD38 transcription which in turn may be associated with improved autistic symptoms^46^.

#### Top-ranked drugs for individual disorders

For SCZ, the top-ranked drug was valproic acid, a widely used mood stabilizer. Although the drug is mainly used for bipolar disorder, meta-analysis of mostly open RCTs showed that it may improve clinical symptoms of SCZ when used as an adjunct to antipsychotics, and may also be useful for aggression or irritability^47^. Minocycline, which is a tetracycline with anti-inflammatory and neuroprotective properties, was ranked among the top and meta-analyses of RCTs have shown its efficacy for SCZ^48,49^. Another drug estriadiol was demonstrated to be effective for treatment-resistant SCZ cases (in women aged 18 to 45) in a double-blind RCT^50^.

For ASD, valproic acid was ranked among the top. Antiepileptics such as valproic acid are frequently prescribed for the control of irritability in patients^51^, although they are not directed for the core symptoms of the disorder. As discussed above, several HDAC inhibitors including trichostatin A and MS-275 were highly ranked.

For ID, several AMPA/kainate receptor antagonists including YM-90K, nbqx and dnqx were highly ranked. As abnormal excitatory glutamatergic transmission is implicated in many neurological and psychiatric disorders, these drugs are being developed for the treatment of a range of diseases, such as epilepsy, cerebral ischemia and Alzheimer disease^52,53^. Several HDAC inhibitors and two antiepileptics (which are also mood stabilizers) valproic acid and carbamazepine were also on the top list.

For epilepsy, many of the top hits are known anticonvulsants or hypnotics/sedatives that enhance GABA transmission. Manually curated descriptions of each drug and their potential therapeutic relevance are listed in Supplementary Tables 1-4.

## DISCUSSION

In this study, we have explored the usefulness of DNMs in guiding drug discovery by looking for overlap of DNMs with drug-related gene-sets. We discovered that neuropsychiatric drug classes were indeed significantly enriched; in particular, antipsychotics and antiepileptics were the *most* strongly enriched drug classes (out of all level 3 ATC classes) for SCZ and epilepsy respectively. By inspecting individual hits, we also uncovered several interesting drug candidates or mechanisms (e.g. HDAC inhibition and retinoid signaling) that might warrant further investigations.

To our knowledge, this is the first study to investigate the usefulness of DNM in guiding drug discovery for neuropsychiatric disorders. The majority of genetic studies in neuropsychiatry aimed at finding new susceptibility genes for specific disorders, however the translational potential of these findings in terms of drug discovery remains largely unexplored. Several related studies are worth mentioning here. A recent study by Ruderfer et al.^54^ found that gene-sets of antipsychotics were enriched for common and rare genetic variations of SCZ, which were derived from a genome-wide association study (GWAS) meta-analysis and a Swedish case-control exome-sequencing study respectively. Interestingly, they also found that agents against amoebiasis and other protozoal diseases was the most significantly enriched drug class^54^. Treatment resistant patients also had an excess of mutations in antipsychotics drug targets^54^. Another study by Gasper et al.^55^ reported that as sample size increases, SCZ GWAS results were increasingly enriched for known psychiatric medications. We also recently revealed that GWAS results of depression and anxiety disorders (and related phenotypes) were enriched for psychiatric drug classes including antidepressants^56^. Taken together, the current study adds to the mounting evidence that data from human genomic studies are useful in guiding drug development in neuropsychiatry.

Most DNM are rare and typically only few patients may share mutations in the same gene. An ultimate goal of drug discovery is to find an effective personalized therapy for *each* genetic subtype of the disorder. This is an important long-term goal but its success might be limited by the very high cost of drug development. An alternative approach is to consider these DNM genes as useful therapeutic targets in general. Here we followed this paradigm with a focus on multi-target^13^ agents. Indeed we found that antipsychotics, which are generally effective for SCZ patients, is the most strongly enriched drug class from DNM. This provides a proof-of-concept of this approach in finding new therapeutic agents. Success stories of drug discovery in the field of cardiovascular medicine also support that rare genetic variations can lead to development of therapies useful for a wider population. For instance, PCSK9 inhibitors are now proven to reduce low-density lipoprotein (LDL) cholesterol and reduce cardiovascular events^57^. The importance of PCSK9 in regulating lipid metabolism was first discovered from rare mutations in familial hypercholesterolemia^58,59^. SGLT2 inhibitor is a new class of anti-diabetic agent^60^; notably rare mutations in the *SGLT2* gene cause familial renal glycosuria^61^.

A few drug classes with significant enrichment are worth mentioning. For SCZ, although antipsychotics ranked first, lipid-lowering agents were also highly significant (*p* = 5.19E-9). This observation is partially supported by clinical studies. For example statins have been shown to ameliorate SCZ symptoms in two small-scale clinical trials^62,63^. In one study simvastatin as adjunctive treatment was tested and there was preliminary evidence for improving total symptoms scores, but the result was not statistically significant^62^. Another study investigated pravastatin also as an adjunctive therapy and found a significant reduction of positive symptoms at week 6, though the effect failed to maintain at week 12^63^. We also found an enrichment of antimicrobial drugs (ATC class code G01A) and antifungal agents (D01A). This is consistent with Ruderfer et al.^54^ who found an enrichment of agents against amoebiasis and other protozoal diseases (P01A); we noted that many drugs such as quinoline derivatives and imidazole derivatives overlap. It should however be noted that this study focused on DNMs, which is different from Ruderfer et al. which investigated GWAS common variants and rare variants from a case-control exome study^54^. Another interesting observation is that antineoplastic agents were ranked among the top for the four neurodevelopmental disorders under study. Several reviews have commented on the possible link between autism and cancer^64,65^. For example, a recent review by Crawley et al.^65^ pointed out several genes and signaling pathways that may be shared between autism and cancer, and suggested that anti-cancer drugs with reasonable safety profiles may be repositioned for ASD. Here we provided for the first time a quantitative assessment of this hypothesis, and indeed revealed enrichment of anti-neoplastic agents.

Another finding from the drug class enrichment analysis is that there is an enrichment of different types of medications (*e.g.* antidepressants/anxiolytics, antiepileptics) across diagnostic categories. Depression and anxiety are common comorbidities for all four disorders under study. There is good evidence to suggest increased rates of depression and anxiety disorders in patients with SCZ^66^ and epilepsy^67,68^. Depressive and anxiety symptoms also appeared to be prevalent in patients with ASD^69,70^ and ID^71-73^. In addition, epilepsy is a well-known comorbidity in ASD and ID^74,75^. The original paper describing the NPdenovo database^4^ demonstrated a significant overlap in DNMs among all four neurodevelopmental disorders, implying a shared pathophysiology. As a result, overlaps in enriched drugs or drug classes are expected.

There are a few limitations to this study. Firstly, we presented an analytic approach to drug discovery or repositioning using DNMs but the study itself did not provide confirmatory evidence for any of the repositioning candidates to be applied in clinical practice. Further pre-clinical and clinical studies, including RCTs, are necessary to verify the efficacy of the drug or drug classes highlighted in this report. In addition, it should be noted that the current gene-set analysis does not take into account the direction of effects. To delineate the direction of drug effects, further work might be needed to characterize the functional impact of mutations as well as how each drug acts on different receptors and genes. Also, before application to clinical practice, the pharmacological properties of each drug, particularly its safety and side-effects, need to be carefully considered. Their permeability through the blood-brain barrier might also affect their effectiveness. These areas have not been investigated in this study. Finally, we mainly focused on DNMs in this work, but further integrative analyses with other sources of genomic data (e.g. GWAS) and drug-related information (e.g. chemical structure and properties), as well as other computational or experimental drug repositioning methods, might improve the chance of discovering effective therapeutics.

In summary, we demonstrated that DNMs might be useful in guiding drug discovery or repositioning. We presented a gene-set analysis approach to achieve this aim and showed that the approach is able to pick up known drugs for the respective disorders; our analysis also highlights a number of repositioning candidates. We hope this work will open a new avenue of finding new therapeutics for neuropsychiatric disorders, and will stimulate further translational research with human genomics data.

## Acknowledgements

This study is partially supported by the Lo-Kwee Seong Biomedical Research Fund and a Direct Grant from the Chinese University of Hong Kong. We would like to thank Mr. Carlos Chau for assistance in data analysis. We would like to thank the Hong Kong Bioinformatics Center for computing support.

## Conflicts of interest

The author declares no conflict of interest.

